# Sparkling Water Consumption Mitigates Cognitive Fatigue during Prolonged Esports Play

**DOI:** 10.1101/2025.06.27.662064

**Authors:** Shion Takahashi, Wataru Kosugi, Seiichi Mizuno, Takashi Matsui

## Abstract

Prolonged esports play induces cognitive fatigue, characterized by diminished executive function with pupil constriction. Players often rely on caffeinated or sugary drinks to combat fatigue, but regular use poses health risks. Sparkling water, a sugar- and caffeine-free beverage, stimulates brainstem and prefrontal activity via sensory pathways mediated by transient receptor potential (TRP) channels in the throat. We here tested the hypothesis that sparkling water mitigates cognitive fatigue during prolonged esports play. Fifteen young adult players participated in a randomized crossover trial, each completing two 3-hour sessions of a virtual football game while consuming either sparkling water or plain water. Subjective fatigue, enjoyment, and executive function (via a flanker task) were measured at baseline and hourly, while pupil diameter and heart rate were monitored continuously. Blood glucose and salivary cortisol were also assessed periodically.

Compared to plain water, sparkling water significantly attenuated increases in subjective fatigue, enhanced enjoyment, and preserved executive function, along with preventing pupil constriction. Heart rate, blood glucose, and salivary cortisol levels did not differ between conditions. Notably, players committed fewer in-game fouls with sparkling water, while offensive and defensive performance remained unchanged. These findings demonstrate that sparkling water alleviates both subjective and objective signs of cognitive fatigue during prolonged esports play, supporting our hypothesis. This non-caffeinated intervention may help sustain inhibitory control and promote fair behavior, offering a safe and sustainable strategy for managing mental fatigue in modern life.

**Highlights:**

- Sparkling water alleviates cognitive fatigue and boosts enjoyment during esports.
- Sparkling water prevents pupil constriction linked to cognitive fatigue.
- Sparkling water keeps players alert without raising blood glucose or cortisol.
- Sparkling water reduces fouls in virtual football without impairing performance.
- Sparkling water offers a caffeine- and sugar-free option to manage mental fatigue.

**Graphical Abstract:** 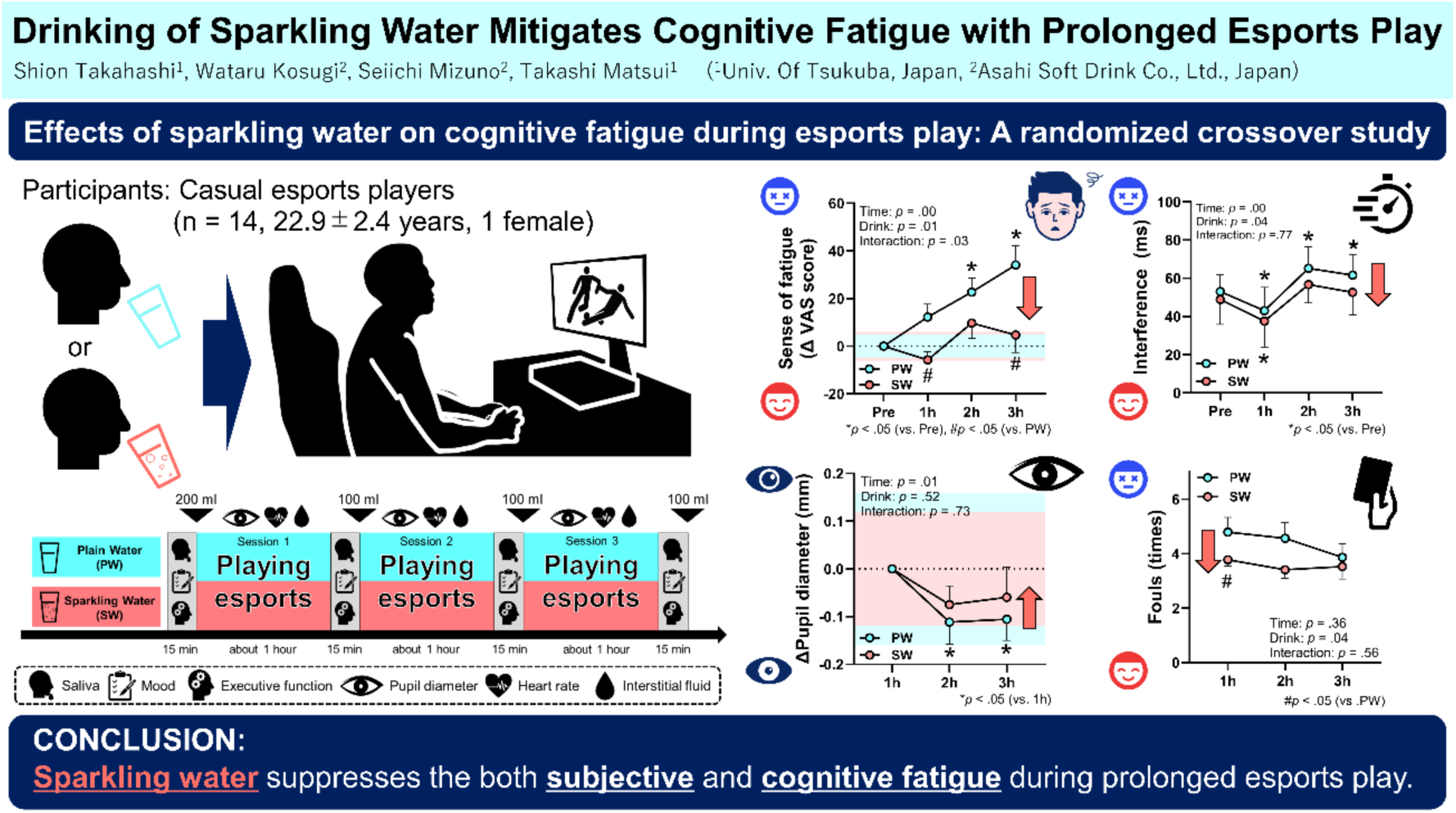

## 1. Introduction

In modern society, people engage in diverse social interactions involving both physical and cognitive activities, which are often accompanied by the accumulation of fatigue (Engberg et al., 2017).

Fatigue can be broadly categorized into functional impairments and the subjective sensation of tiredness. The sensation of fatigue, in particular, serves as a biological defense mechanism that temporarily limits activity levels to prevent excessive exertion, functioning similarly to pain (Gibson et al., 2003). However, during mentally demanding activities that do not involve physical movement—such as visual display terminal (VDT) work or prolonged driving—cognitive functions may deteriorate without a concurrent sense of fatigue (Sun et al., 2022; Schmidt et al., 2009), a condition referred to as cognitive fatigue (Pessiglione et al., 2025). For modern humans, whose lifestyles increasingly separate physical from mental activity, addressing cognitive fatigue could help prevent severe accidents and long-term health problems associated with unnoticed cognitive decline (Sasahara et al., 2015).

Esports exemplify a contemporary form of digital cognitive engagement. These competitive video game-based activities have expanded rapidly in recent years, particularly among younger populations (Jenny et al., 2017). While physical ability is central in traditional sports, esports rely on digital interaction, requiring advanced cognitive skills such as spatial reasoning, sustained attention, information processing, and task switching (Miao et al., 2024; Toth et al., 2020). Indeed, prolonged esports play in such cognitively demanding environments reduces executive function and pupil diameter, a neurobiomarker of prefrontal activity, even before the onset of subjective fatigue, regardless of a player’s level of expertise (Matsui et al., 2024). Therefore, understanding the mechanisms of cognitive fatigue in esports and identifying effective countermeasures are essential for managing both performance and health of the body and mind in modern population.

To combat fatigue, modern people including esports players frequently consume beverages such as energy drinks and coffee (Pang et al., 2025). These drinks contain caffeine and glucose, which enhance executive function upon ingestion (Sainz et al., 2020; Wu et al., 2024), thereby delaying the progression of cognitive fatigue (Kennedy & Scholey, 2004). However, excessive or long-term intake of caffeine and sugar is associated with adverse health effects, including abnormal heart rate (Berger & Alford, 2009), sleep disturbances (Machado-Vieira et al., 2001), impaired glucose tolerance (Bedi et al., 2014), and increased depressive symptoms (Kaur et al., 2020). These concerns highlight the need for safer and more sustainable beverage alternatives to help manage cognitive fatigue.

Sparkling water, an unsweetened carbonated beverage free of caffeine and sugar, has gained attention as a health-conscious option. Regular consumption of sparkling water has become more common in recent years (Grand View Research, 2021). A previous study showed that sparkling water consumption improves mood by inducing a sense of exhilaration, enhancing motivation, and reducing sleepiness (Fujii et al., 2022). Moreover, drinking carbonated beverages delays the onset of subjective fatigue associated with cognitive activity (Smit et al., 2004). Thus, sparkling water could help suppress subjective fatigue during esports.

In addition to its mood-related effects, sparkling water may also influence cognitive function directly. Carbonation in energy drinks enhances executive performance in simple cognitive tasks (Smit et al., 2004). These effects are likely mediated by carbon dioxide (CO₂), which stimulates TRPA1 channels in the trigeminal nerve (Wang et al., 2010). CO₂ released as bubbles, a characteristic feature of sparkling water, also activates TRPV1 channels in the pharynx and brainstem, key components of the ascending reticular activating system (Tsuji et al., 2020). Also, sparkling water activates regions of the prefrontal cortex involved in executive function, such as the orbitofrontal cortex (OFC), likely via stimulation of this ascending arousal system (Zhang et al., 2020; Kosugi et al., 2024). Therefore, sparkling water could activate prefrontal cortex and mitigate cognitive decline associated with prolonged esports play.

We thus hypothesized that drinking sparkling water suppresses both subjective fatigue and cognitive fatigue during prolonged esports play. To test this hypothesis, we conducted a randomized crossover experiment in which young adult players who regularly engage in video gaming were instructed to play a virtual soccer game for three hours while consuming either plain (noncarbonated) or sparkling water. To evaluate the effects of sparkling water on cognitive fatigue, we measured cognitive and neurophysiological indices during esports play.

## 2. Material and Method

### 2.1. Participants

We recruited 15 casual players from the student population at the University of Tsukuba who reported regularly playing video games. The definition of a casual player was based on the criteria used in Matsui et al. (2024). Of these, 14 participants (22.9 ± 2.4 years old; 1 female) were included in the final analysis, with one excluded due to data recording errors (Table 1). Eligibility criteria included: age between 18 and 35 years, capacity to understand and sign informed consent, normal color vision, and availability to attend the experiment in person. Ethical approval for this study was granted by the Research Ethics Board of the University of Tsukuba, and the procedures complied with the Declaration of Helsinki.

### 2.2 Pre-experiment questionnaire

Following the method described by Monma et al. (2024), demographic data including esports playing habits, physical activity levels, and sleep duration were collected using a Google Forms questionnaire. Participants received a survey link and study outline via email at the time of recruitment and were instructed to complete the form by the morning of the day before the measurement. Esports status was determined by self-reported main title played, duration of experience (months), rank, and average playtime (minutes) on weekdays and weekends. Physical activity was assessed using the Japanese version of the International Physical Activity Questionnaire (IPAQ), as validated by Murase et al. (2002).

**Table 1.** Participant characteristics of playing lifestyles.

|  |  | n = 14 |  |
| --- | --- | --- | --- |
| Age | Years | 22.9 | ± 2.4 |
| Sex | Male/female | 13 | / 1 |
| Weight | kg | 69.4 | ± 15.2 |
| Height | m | 1.7 | ± 0.1 |
| BMI | kg/m<sup>2</sup> | 23.5 | ± 3.5 |
| Playing career of<br>proficient title | months | 5.5 | ± 5.3 |
| In-game rank | top percentile | 66.9 | ± 25.5 |
| Weekday esports<br>playing time | min/day | 65.5 | ± 74.5 |
| Weekend esports<br>playing time | min/day | 114.5 | ± 115.3 |
| Sleeping time | min/day | 425.5 | ± 18.1 |
| Vigorous-intensity<br>activity time | min/day | 64.1 | ± 78.3 |
| Moderate-intensity<br>activity time | min/day | 81.8 | ± 116.6 |
| Walking time | min/day | 23.2 | ± 17.9 |
| Exercise time | min/day | 27.3 | ± 19.7 |
| Sitting time | min/day | 387.3 | ± 263.4 |

Data are shown as mean ± SEM. Sex data are presented as the number of male and female participants. The in-game rank data show percentages from the highest to the lowest rank, ranging from 0 to 100.

### 2.3. Body composition

We assessed participants’ body composition using a bioelectrical impedance analyzer (Seca mBCA 515; Seca, Hamburg, Germany), a device frequently used in esports research (Matsui et al., 2024; Krarup et al., 2024). Participants stood barefoot on the device while four electrodes were placed on the limbs, followed by a waist circumference measurement. Parameters recorded included height, weight, BMI, fat mass, regional skeletal muscle mass, body water content, and energy expenditure. All measurements were conducted 2–7 days before the experiment.

### 2.4. Experimental protocol

The experimental procedure followed the method previously validated by Matsui et al. (2024) for assessing cognitive fatigue in esports players. Two to seven days prior to the experiment, participants played virtual football (eFootball, Konami Digital Entertainment Co., Ltd., Japan) for at least one hour at the measurement site to acclimate to the game environment and cognitive tasks.

Participants were instructed to refrain from consuming alcohol from the day before until the end of the experiment, from consuming caffeine and engaging in strenuous physical activity on the day of the experiment, and from eating during the two hours preceding the session.

The experimental protocol is illustrated in Figure 1. On the day of measurement, the laboratory was maintained at a temperature of 24 ± 2°C and a humidity of 50 ± 10%. Upon arrival, participants received a verbal explanation of the experimental procedures and provided written informed consent. A heart rate monitor (Polar H10; Polar, Finland) was attached to the chest, after which participants rested in a seated position for 10 minutes. Following the rest period, participants played a virtual soccer game against a standard-level CPU for a total duration of three hours.

**Figure 1.**
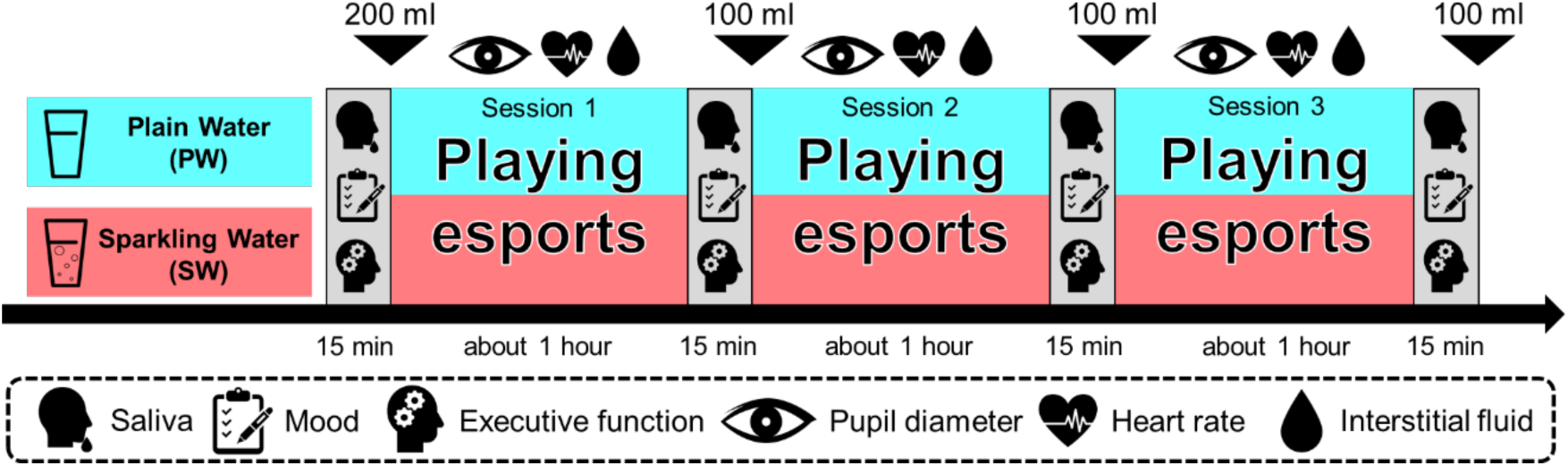
Protocol of experiments. The same protocol was applied to the plain water (PW) and sparkling water (SW). Participants engaged in eFootball gameplay against a standard-level CPU for a total duration of three hours. Saliva sample collection, administration of subjective mood questionnaires, and executive function assessments were conducted before the session commenced and at hourly intervals throughout the gameplay. Each assessment session lasted approximately 15 minutes. During gameplay, pupil diameter and heart rate were continuously monitored. At the time points indicated by the downward- facing triangulator in the figure, participants consumed either plain or sparkling water. The beverages were chilled to 4°C, poured into paper cups, and provided to participants immediately before consumption.

Prior to each play session, participants consumed either plain or sparkling water, chilled to 4°C, within approximately two minutes. The volume was 200 ml before the first session and 100 ml before each of the following two sessions, totaling 500 ml. All beverages were stored in a refrigerator, opened just before consumption, and served in disposable paper cups.

Subjective sensations and executive function were assessed before gameplay (Pre) and at one-hour intervals during the session (1 h, 2 h, and 3 h). Saliva samples were collected at the same time points to measure cortisol levels. Throughout gameplay, pupil diameter was continuously recorded using an eye tracker (Tobii Pro Nano; Tobii Technology Co., Ltd., Sweden). Interstitial fluid glucose levels were monitored continuously throughout this experiment using the FreeStyle Libre Pro (Abbott Diabetes Care, Witney, UK).

At least two days later, participants completed the same procedure with the alternative beverage. Each session lasted approximately 300 minutes, from arrival to dismissal. All participants received an honorarium of 5,000 yen upon completion.

### 2.5. Questionnaires

Participants rated two subjective sensations, enjoyment and fatigue, using a 100-mm visual analog scale (VAS), in accordance with previous studies (Tseng et al., 2010; Kotz et al., 2011). The VAS was a horizontal line anchored with opposing phrases at each end: “enjoyed it not at all” to “enjoyed it very much” for enjoyment, and “no fatigue” to “very severe fatigue” for fatigue. Participants marked the line at the point that best represented their current state. The distance from the left end (in mm) was measured and converted into a percentage of the total line length.

### 2.6. Executive functions

Executive function was assessed during and after esports play using flanker task, with PsychoPy3 (v2021.1.2) according to a task condition used in our previous study (Matsui et al. 2024). Stimuli were presented in random order with an equal number of congruent and incongruent trials. Each trial consisted of a fixation point (250 ms), stimulus presentation (up to 2000 ms), and a blank screen (1000 ms). The stimulus disappeared upon keypress and was followed by a blank screen. Reaction times and accuracy were recorded. The interference effect was calculated as the difference in reaction time between incongruent and congruent trials.

### 2.7. Pupillometry

Following the protocol described by Matsui et al. (2024), pupil diameter was recorded as an index of prefrontal activity using an infrared eye tracker (Tobii Pro Nano; Tobii Technology, Sweden) during esports play. Ambient lighting was standardized to 250–300 lux at eye level. The eye tracker was positioned below the game display and connected to a dedicated laptop. Participants were seated approximately 60 cm from the screen. Calibration was performed before the first session, and recordings were taken continuously throughout all three sessions at a sampling rate of 60 Hz. Only successfully recorded data segments were included in the analysis.

### 2.8. Related physiological measurements

To assess physiological responses to prolonged esports play, we measured heart rate, interstitial fluid glucose, and salivary cortisol, following prior studies in esports players (Matsui et al., 2024; Cregan et al., 2025). Heart rate was recorded continuously at 1 Hz using a chest-worn band-type monitor (Polar H10; Polar, Finland) worn from the beginning of the experimental session.

Interstitial fluid glucose was used as a non-invasive proxy for blood glucose variability. A continuous glucose monitor (FreeStyle Libre Pro; Abbott Diabetes Care, Witney, UK) was attached to the back of the upper arm 2–7 days before the main experiment. Glucose levels were recorded at 15-minute intervals. Salivary cortisol was measured as a biomarker of physiological stress. Saliva samples (2 ml each) were collected via straw, immediately frozen at −80 °C, and later centrifuged at 1,500 × g for 20 minutes to remove mucus. The resulting supernatant was aliquoted and stored at −80 °C until analysis. Cortisol concentrations were quantified in duplicate using a commercial ELISA kit (Salivary Cortisol ELISA Kit; Salimetrics, LLC.), following the manufacturer’s instructions. Absorbance was measured, and cortisol concentrations were calculated using standard curves generated with a microplate reader (Varioskan LUX Multimode Microplate Reader, Thermo Fisher Scientific, MA).

### 2.9. Statistical analysis

All statistical analyses were conducted using GraphPad Prism version 10 (GraphPad Software, San Diego, CA). Data were analyzed using either two-way analysis of variance (ANOVA) or paired t-tests, depending on the comparison. To assess the relationship between pupil diameter and executive function, we conducted Pearson’s correlation analysis. When significant main effects or interactions were found, Bonferroni’s post hoc comparisons were performed. Results are presented as means ± standard error of the mean (SEM). A p-value < 0.05 was considered statistically significant.

## 3. Results

### 3.1. Drinking sparkling water suppresses the sense of fatigue and enhances the enjoyment

To assess how players feel when drinking sparkling water during esports play, we first evaluated their subjective sensations, sense of fatigue and enjoyment, using a visual analog scale (VAS) (Figure 2). No significant differences were found in baseline VAS scores between conditions (Table 2). In the plain water condition, fatigue increased significantly after more than two hours of gameplay, whereas it did not increase in the sparkling water condition (Figure 2A). Enjoyment increased significantly over time in both conditions, with a greater increase observed in the sparkling water condition (Figure 2B). These results suggest that drinking sparkling water suppresses the development of subjective fatigue and enhances enjoyment during prolonged esports play.

**Figure 2.**
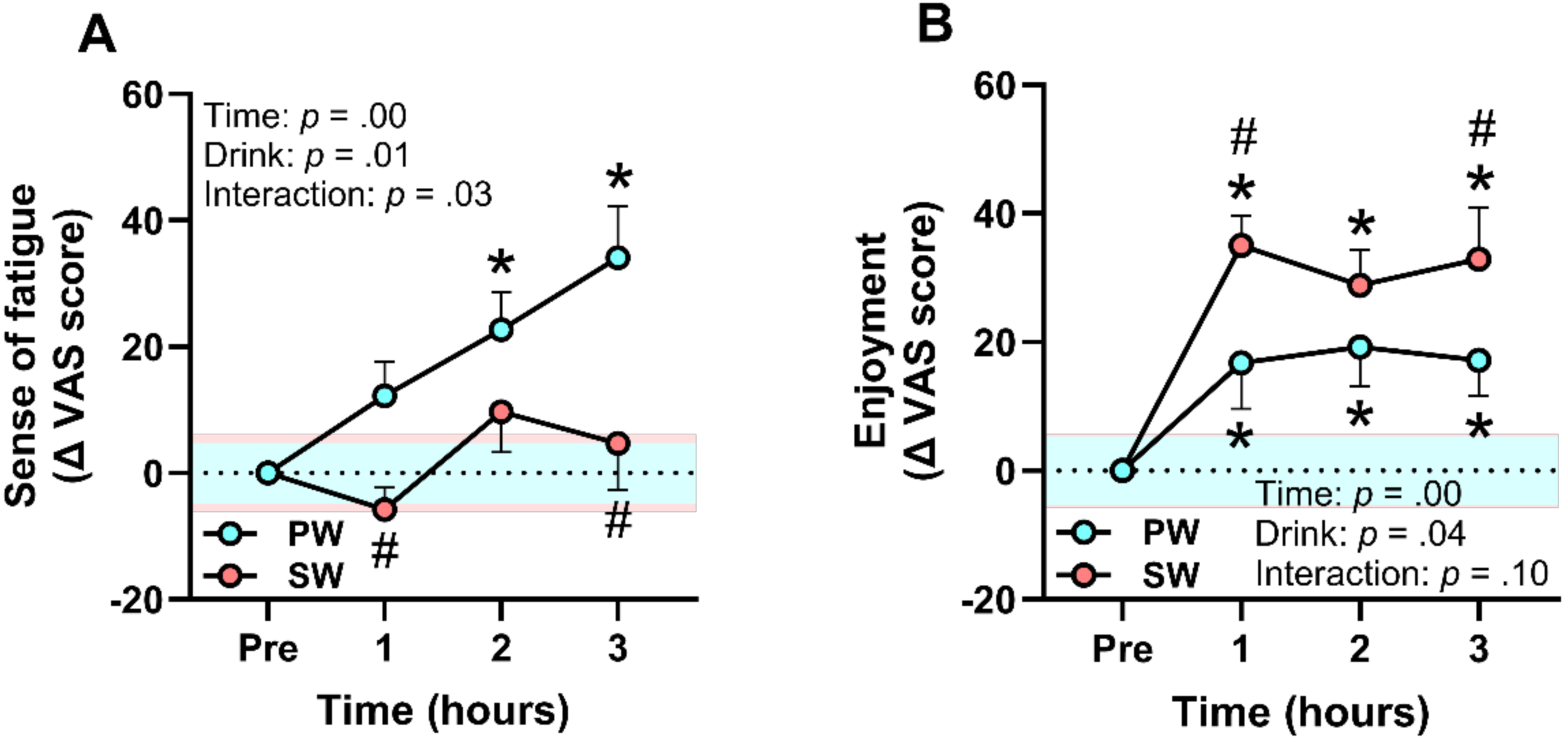
Sparkling water consumption inhibits the onset of a sense of fatigue and enhances enjoyment during prolonged esports play. **A.** Amount of change in the VAS score for the sense of fatigue from Pre-point. **B.** Amount of change in the VAS score for the enjoyment from Pre-point. Data are shown as mean ± SEM. The blue and red bands in both graphs represent the SEM of the absolute VAS score at the Pre-point for PW and CW, respectively. Results of two-way ANOVA are shown in the corner of each graph. *Time*: main effect of playing time. *Drink*: main effect of drink condition. *Interaction*: interaction between playing time and drink condition. \**p* < .05 vs. Pre for both groups; *close proximity to a point on the graph. #*p* < .05 vs. PW. The results representing * and # were calculated using Bonferroni’s multiple comparison test.

**Table 2.** Psychophysiological parameters at base line (Pre)

|  |  | Plain Water<br>(n = 15) |  |  | Sparkling Water<br>(n = 15) |  |  |
| --- | --- | --- | --- | --- | --- | --- | --- |
| Sense of fatigue | VAS score | 26.4 | ± | 5.3 | 33.7 | ± | 6.5 |
| Enjoyment | VAS score | 40.3 | ± | 5.7 | 32.6 | ± | 5.1 |
| Pupil diameter | mm | 4.0 | ± | 0.2 | 4.0 | ± | 0.1 |
| Heart rate | bpm | 79.1 | ± | 4.8 | 82.8 | ± | 3.7 |
| Glucose levels | mg/dL | 88.3 | ± | 5.9 | 91.2 | ± | 3.3 |
| Cortisol levels | µg/dL | 0.2 | ± | 0.0 | 0.2 | ± | 0.0 |

Data are expressed as mean ± SEM. Statistical differences were calculated by paired t-test. There were no significant differences in these items.

### 3.2. Sparkling water prevents cognitive fatigue induced by prolonged esports play

To evaluate the anti-cognitive fatigue effect of sparkling water, we assessed the dynamics of executive function using the flanker task. In the plain water condition, cognitive fatigue was evident as an increase in flanker interference time after more than two hours of play (Figure 3A), and a decrease in the percentage of correct responses to incongruent trials after three hours (Figure 3B). In contrast, the sparkling water condition preserved the improvement in interference time observed at one hour and suppressed the increase observed after two or more hours of play (Figure 3A).

**Figure 3.**
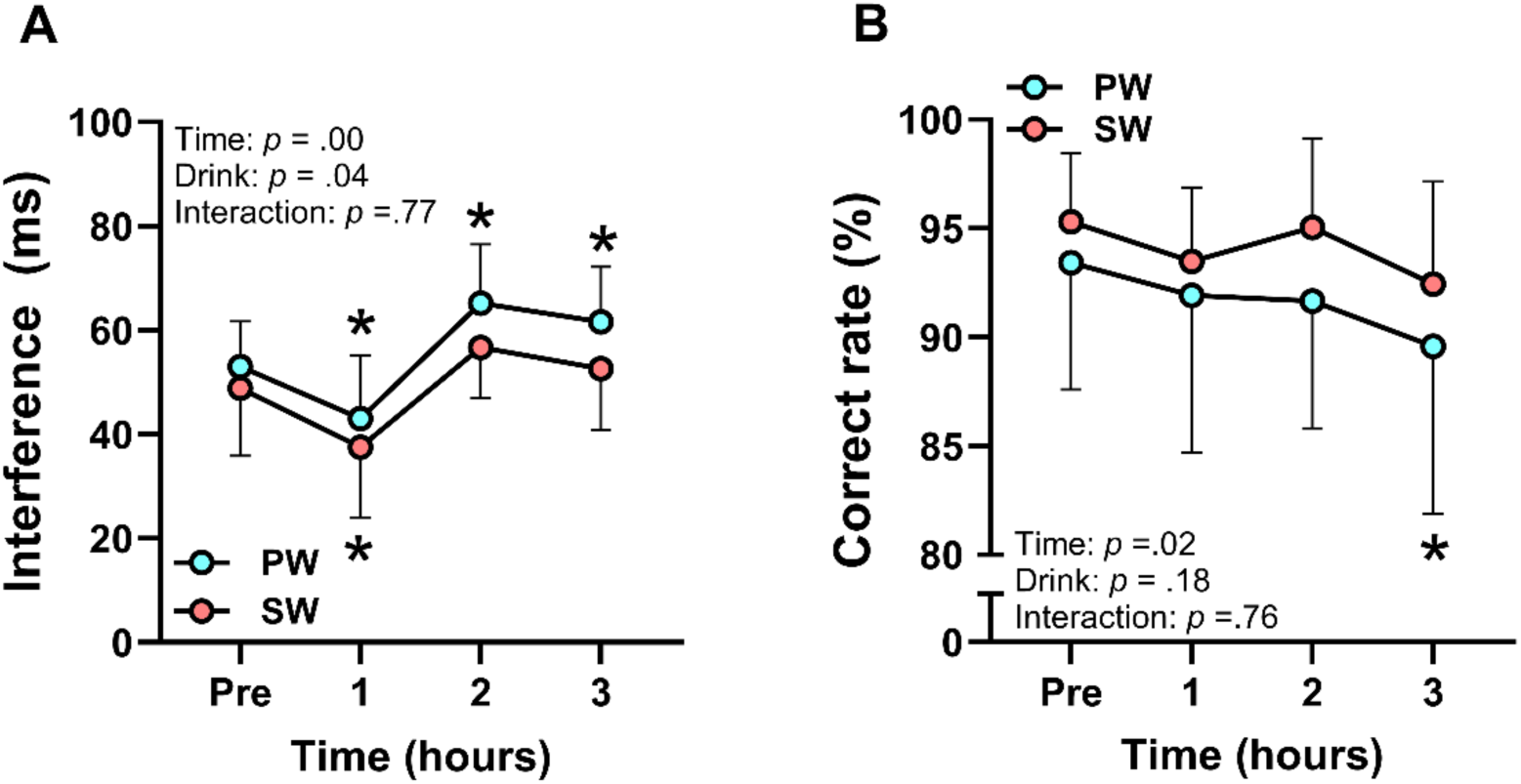
Drinking sparkling water helps to maintain cognitive performance in terms of both speed and accuracy during prolonged esports play. **A.** Interference time in the flanker task. **B.** Percentage of correct responses in the incongruent flanker task. Data are shown as mean ± SEM. Results of two-way ANOVA are shown in the corner of each graph. *Time*: main effect of playing time. *Drink*: main effect of drink condition. *Interaction*: interaction between playing time and drink condition. \**p* < .05 vs. Pre for both groups; *close proximity to a point on the graph. #*p* < .05 vs. PW. The results representing * and # were calculated using Bonferroni’s multiple comparison test.

Furthermore, the correct response rate in the incongruent task did not decline over the course of the experiment (Figure 3B). These results suggest that sparkling water consumption prevents cognitive fatigue during prolonged esports play.

### 3.3. Sparkling water attenuates the reduction in pupil diameter, independently of blood glucose and stress response levels

To investigate the physiological mechanisms underlying cognitive fatigue-reducing effect of sparkling water, we assessed non-invasive physiological indices that did not interfere with play, including pupil diameter. Baseline values of all physiological indices did not differ significantly between conditions (Table 2). In the plain water condition, pupil diameter decreased after the first two hours of play, but no such reduction was observed in the sparkling water condition (Figure 4A). Neither average heart rate nor maximum heart rate changed over the course of the experiment (Supplementary Figure 1, Figure 4B). Interstitial fluid glucose levels gradually decreased over time, but no beverage-related effects were observed (Figure 4C). Salivary cortisol levels remained unchanged regardless of beverage condition (Figure 4D). These findings suggest that sparkling water suppresses the reduction in pupil diameter as an indicator of prefrontal activity, independently of changes in energy metabolism or stress levels during prolonged esports play.

**Figure 4.**
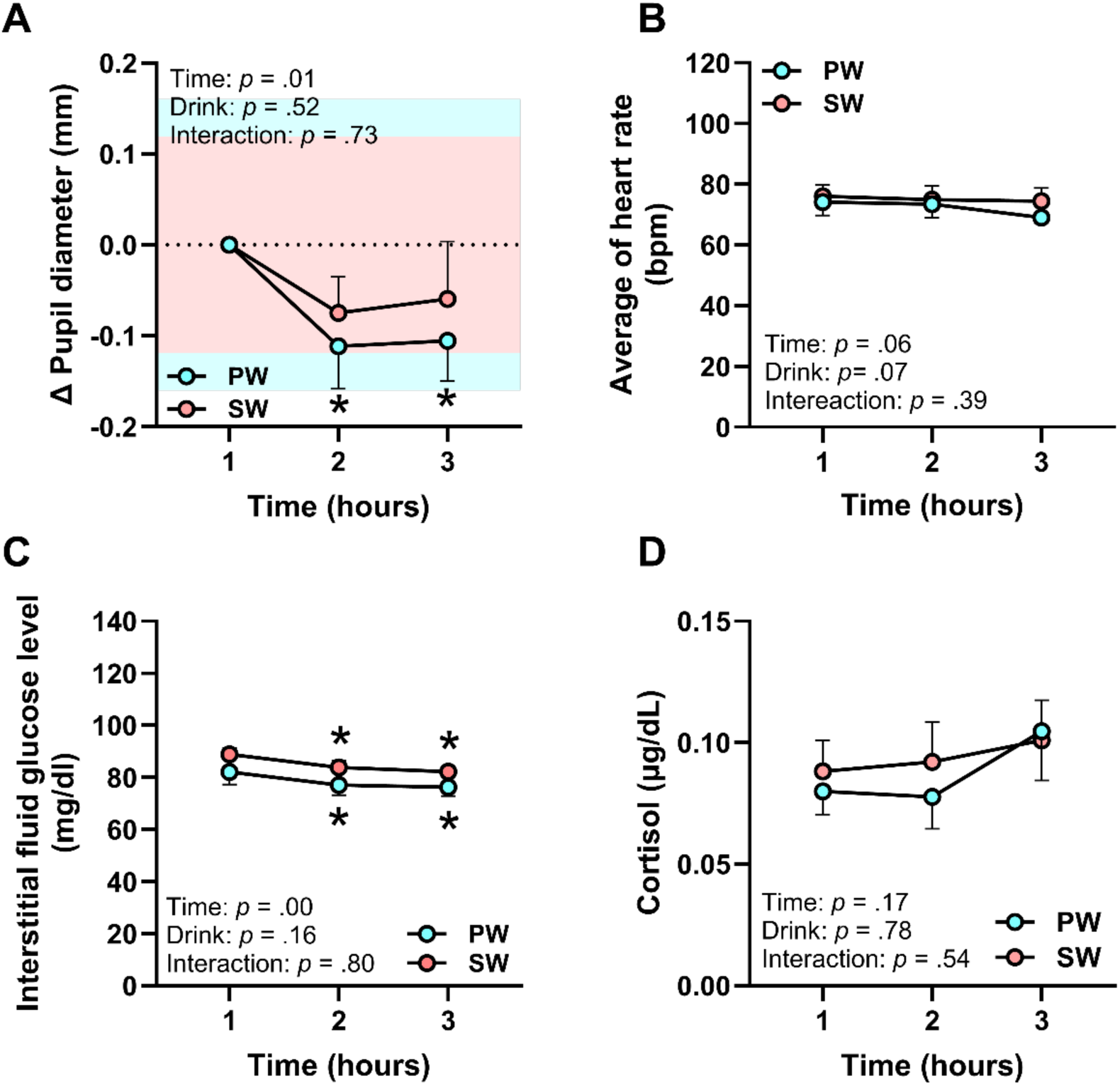
Drinking sparkling water without providing energy can suppress pupil diameter reduction and increase maximal heart rate during prolonged esports. **A.** The amount of change of pupil diameter from 1 h. **B.** Maximum hourly heart rate of participants during esports play. **C.** Interstitial fluid glucose levels of participants throughout the experiment. **D.** Trends in hourly salivary cortisol levels of participants. Data are shown as mean ± SEM. The blue and red bands in A represent the SEM of the absolute pupil diameter at 1h for plain water (PW) and sparkling water (SW), respectively. Results of two-way ANOVA are shown in the lower-left corner of each graph. *Time*: main effect of playing time. *Drink*: main effect of drink condition. *Interaction*: interaction between playing time and drink condition. \**p* < .05 vs. 1 h for both groups; *close proximity to a point on the graph. #*p* < .05 vs. PW. The results representing * and # were calculated using Bonferroni’s multiple comparison test.

### 3.4. Maintenance of pupil diameter is associated with suppressed cognitive fatigue

To determine whether the anti-cognitive fatigue effect of sparkling water is associated with changes in pupil diameter, we analyzed the correlation between flanker interference time and pupil diameter change. When data from both conditions were integrated, a significant negative correlation was observed between the two indices (Figure 5A). When the correlations were examined separately by the beverage condition, we observed a significant correlation in plain water condition but no significant correlation was found in the sparkling water condition (Figure 5B, C). These results suggest that sparkling water consumption mitigates cognitive fatigue by preventing reductions in prefrontal activation, as indexed by pupil size.

**Figure 5.**
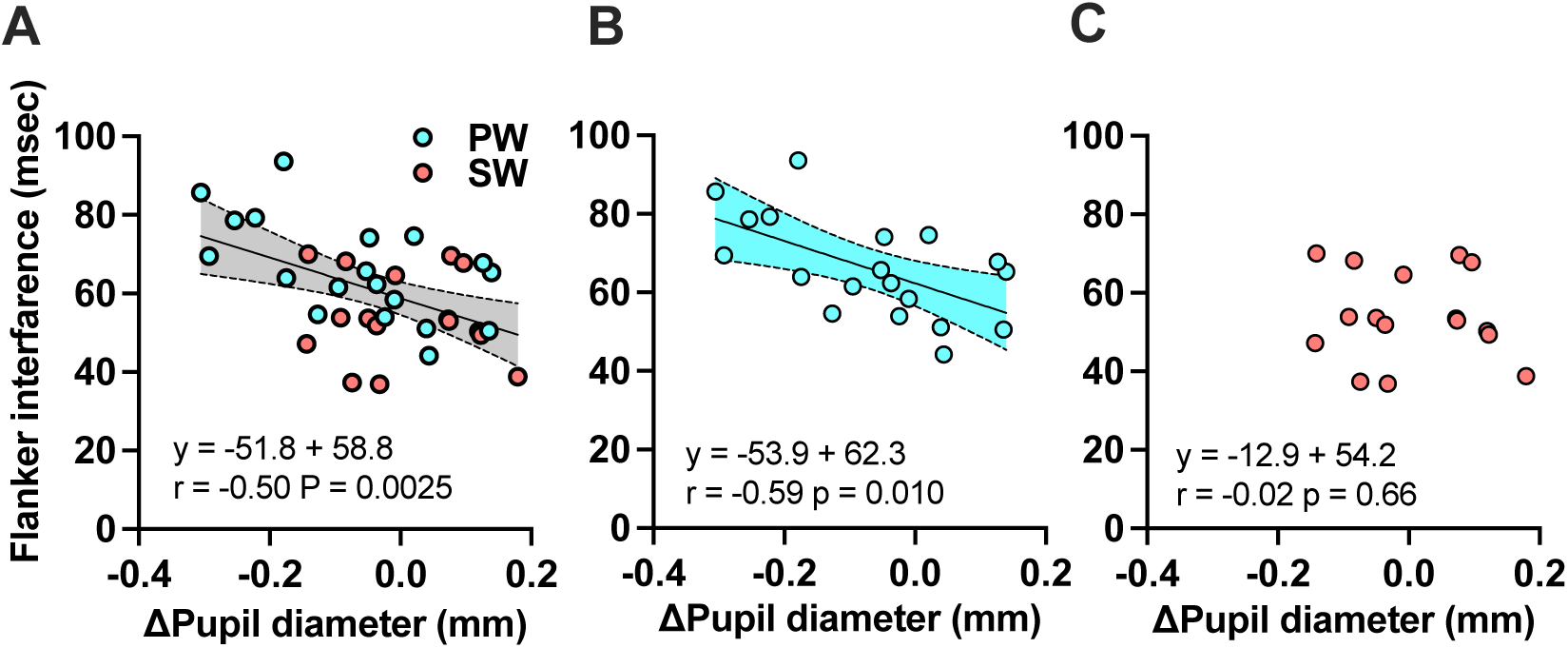
Cognitive decline is associated with pupil constriction. Correlation between pupil diameter change and flanker interference time including both drink condition (A), in plain water (PW, B) and in sparkling water (SW, C). The Pearson’s product-rate correlation coefficient and the results of the single regression analysis are shown in the lower-left corner of the graph. The computed regression line and 95% confidence intervals are shown as a gray band on the graph.

### 3.5. Drinking sparkling water inhibits the number of fouls during play

To evaluate the effect of sparkling water consumption on in-game performances, average game statistics were analyzed across each one-hour session. Beverage type had no significant effect on key offensive metrics such as the number of passes or shots, which are factors directly related to match outcomes (Figure 6A-E, Supplementary Figure 2). However, the average number of fouls was significantly lower in the sparkling water condition (Figure 6F). These findings suggest that drinking sparkling water during esports play promotes fair play behavior without impairing competitive performance.

**Figure 6.**
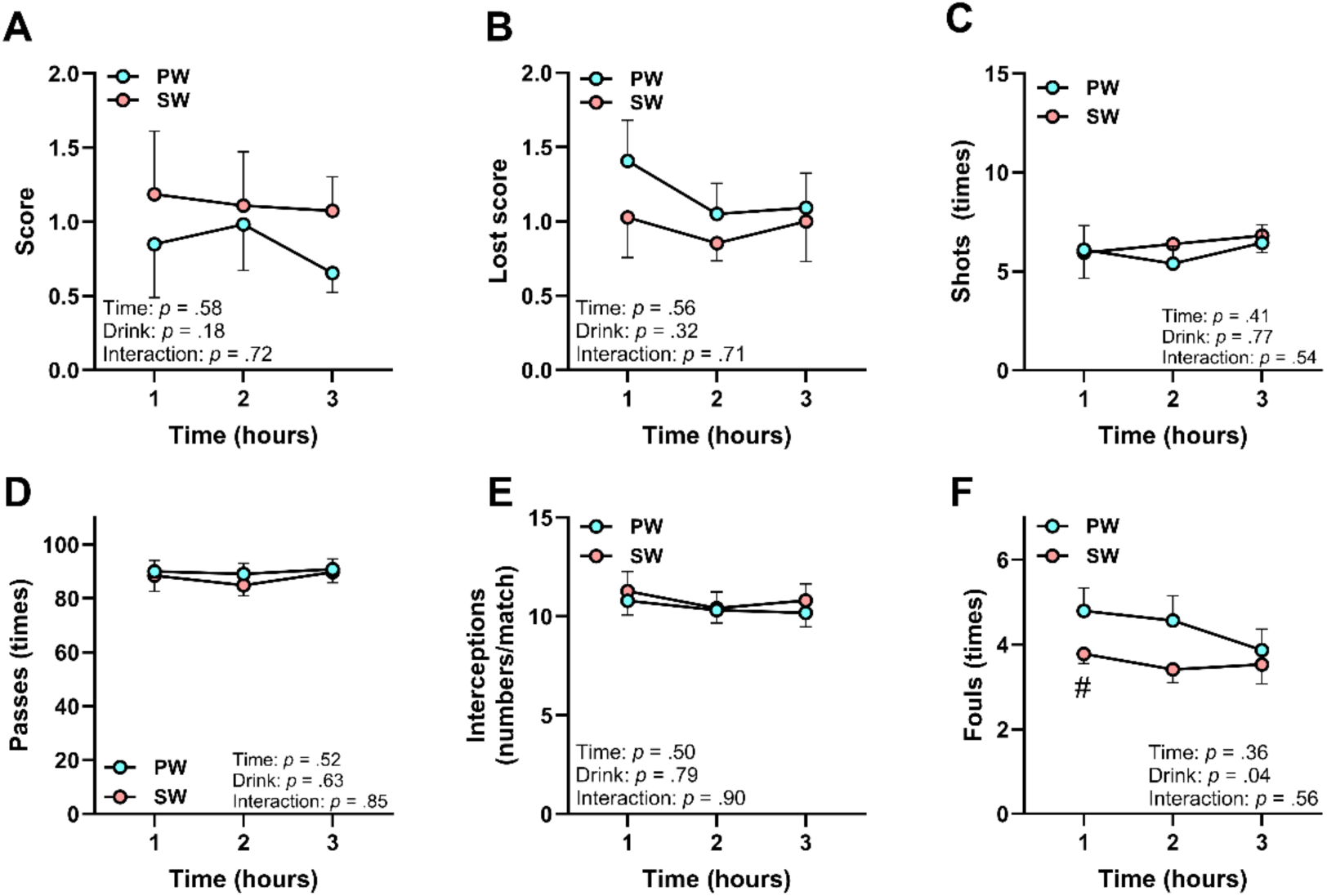
Drinking sparkling water inhibits non-sporting behavior during esports play. **A. S**core. **B.** Score of opponents. **C.** Number of shots. **D.** Number of passes. **E.** Number of interceptions. **F.** Number of fouls. All data are the average per hour. Data are shown as mean ± SEM. Results of two-way ANOVA are shown in the corner of each graph. *Time*: main effect of playing time. *Drink*: main effect of drink condition. *Interaction*: interaction between playing time and drink condition. #*p* < .05 vs. PW. The results representing # was calculated using Bonferroni’s multiple comparison test.

## 4. Discussion

This study tested the hypothesis that drinking sparkling water mitigates both subjective and cognitive fatigue via prefrontal activation during prolonged esports play. We found that sparkling water consumption reduced subjective fatigue, prevented cognitive decline, and attenuated pupil constriction as an indicator of prefrontal activity, supporting our hypothesis. Sparkling water also led to fewer fouls committed during virtual soccer play. Notably, these effects occurred independently of interstitial glucose levels or stress-related physiological responses.

### 4.1. Why are both subjective and cognitive fatigue suppressed?

In the present study, drinking sparkling water suppressed the onset of subjective fatigue during prolonged esports play (Figure 2A). One plausible explanation is that cognitive fatigue itself was suppressed in the sparkling water condition. Although subjective fatigue typically emerges later than cognitive decline during prolonged esports play (Matsui et al., 2024), no signs of cognitive impairment were observed after three hours of esports play in the sparkling water condition (Figure 3A, B). Therefore, it is reasonable to assume that suppressing cognitive fatigue helped prevent the subsequent development of subjective fatigue.

Interestingly, no increase in subjective fatigue was reported in the sparkling water condition even during the early phase of esports play, when cognitive fatigue was not yet present in the plain water condition (Figure 2A). This suggests that the enhanced enjoyment associated with sparkling water contributes to this outcome. Subjective fatigue during task engagement can be masked or perceived as less intense when the task is highly enjoyable (Milyavskaya et al., 2021).

While participants reported enjoyment under both beverage conditions, enjoyment levels were notably higher in the sparkling water condition (Figure 2B). Previous studies showed that sparkling water enhances mood by promoting feelings of exhilaration and motivation (Fujii et al., 2022), and that higher motivation during cognitive tasks is associated with greater enjoyment (Milyavskaya et al., 2021). Therefore, the psychological effects of enhanced enjoyment likely contributes to masking the perception of subjective fatigue during esports play in the sparkling water condition..

### 4.2. Psychophysiological mechanisms of sparkling water in mitigating cognitive fatigue

What neuronal mechanisms underlie the suppression of cognitive fatigue by sparkling water? Pupil diameter may offer an important clue. Pupil size is a neurobiological marker of the ascending arousal system and reflects prefrontal activity, which is modulated by noradrenergic and dopaminergic pathways (Pfeffer et al., 2022). Our previous study showed that pupil constriction is more closely linked to the onset of cognitive fatigue than to subjective fatigue (Matsui et al., 2024). In the present study, pupil diameter remained stable in the sparkling water condition but decreased in the plain water condition during prolonged play (Figure 4A). Additionally, across both beverage conditions, pupil diameter was significantly negatively correlated with flanker task performance (Figure 5A).

These results suggest that sparkling water consumption helps sustain prefrontal activity via ascending arousal pathways.

Carbon dioxide (CO₂) in carbonated beverages activates TRP channels, such as TRPA1 and TRPV1 (Wang et al., 2010; Tsuji et al., 2020). In animal models, activation of these channels has been shown to stimulate brain regions involved in cognitive control, including the brainstem and prefrontal cortex (Tsuji et al., 2020; Zhang et al., 2020). In humans, sparkling water consumption increases local cerebral blood flow in the prefrontal cortex as a center of executive function, particularly in the orbitofrontal cortex (OFC) (Kosugi et al., 2024). Therefore, sparkling water could suppress the onset of cognitive fatigue by maintaining the prefrontal cortical activity through TRP channel–mediated activation.

### 4.3. Why does drinking sparkling water reduce the number of fouls?

Although sparkling water had little effect on most gameplay statistics in virtual football (Figure 6 A-E), it significantly reduced the number of fouls committed (Figure 6F). This suggests that sparkling water promotes fair play without impairing playing performance. One likely mechanism is the prevention of executive function. In eFootball, fouls, whether intentional or unintentional, are assessed according to the real-world soccer rules. Executive functions such as inhibitory control and cognitive flexibility are essential for regulating aggressive behavior and limiting excessive contact, and a decline in these functions contributes to the occurrence of fouls (Mergan, 2022). A previous study also reported that fouls, particularly those resulting in yellow cards, increase in the latter stages of a match as players become fatigued (Sun et al., 2024). Thus, the reduction in fouls observed in the sparkling water condition may reflect the preservation of executive control throughout esports play.

Another possible mechanism is the modulation of negative emotional responses.

Emotional states that can lead to fouls are often negative in valence. The ventromedial prefrontal cortex (vmPFC) is involved in regulating these emotional control (Hiser & Koenigs, 2018). Previous study showed that sparkling water increases local cerebral blood flow in the OFC, which is anatomically close to the vmPFC (Kosugi et al., 2024). Consistent with this, we also indirectly observed that enhanced prefrontal activity in the sparkling water condition, as indicated by pupillometry data (Figure 4A). Taken together, sparkling water may reduce foul behavior by suppressing negative emotions, possibly via vmPFC activation.

### 4.4. Sparkling water as an alternative to caffeine and sugar

Our findings indicate that sparkling water is effective in reducing cognitive fatigue associated with prolonged mental activity, extending beyond the context of esports. Caffeine and sugar are commonly used to counteract cognitive fatigue (Kennedy & Scholey, 2004), and they are typically suited for short-term, high-intensity situations that require elevated focus and rapid response, such as examinations or competitive events (Sainz et al., 2020; Wu et al., 2024). However, chronic use of these substances carries risks due to their potential for dependence and overconsumption, which can lead to adverse physical and mental health outcomes (Bedi et al., 2014; Berger & Alford, 2009; Kaur et al., 2020; Machado-Vieira et al., 2001). In contrast, sparkling water, being free of caffeine and sugar, can be consumed regularly without associated health risks. Therefore, sparkling water should serve as a safer and more sustainable alternative to stimulant-based interventions for managing daily cognitive fatigue.

As digital technology becomes increasingly integrated into modern life, cognitively demanding tasks are becoming routine across domains such as work, education, and healthcare. These tasks often require sustained cognitive effort and performance. Esports, as examined in this study, serves as a representative model of such cognitively intensive activities. Accordingly, the observed anti-cognitive fatigue effects of sparkling water could have broader applicability, potentially benefiting individuals engaged in prolonged mental work, such as office-based tasks or academic study.

### 4.5. Limitations

This study has several limitations. First, the participant sample consisted exclusively of young, casual esports players, which limits the generalizability of the findings. Cognitive fatigue during prolonged esports play is observed across different player types, but variations in gaming expertise affects its onset and expression (Matsui et al., 2024). Future research should include more diverse populations, such as athletically oriented players, to assess whether cognitive fatigue–reducing effects of sparkling water extend across different user profiles. Second, the physiological mechanism underlying the observed effects of sparkling water was not directly examined. While we discussed that carbonation-induced stimulation of pharyngeal TRP channels contributed to the effect, this was not examined directly. Investigating this would require foundational studies—distinct from applied research in esports contexts—using approaches such as pharmacological interventions or TRP knockout animal models. Third, full blinding of beverage conditions was not feasible. Although white opaque cups were used to conceal visual characteristics, the sensory cues, such as carbonation- induced oral irritation, likely allowed participants to identify the beverages. However, participants were unaware of the study’s hypotheses, and the influence of condition awareness was considered minimal. Furthermore, in real-world esports settings, concealing beverage types is often impractical.

Therefore, despite this limitation, the study retains strong ecological validity and relevance to real-life cognitive activities.

## 5. Conclusion

Our findings demonstrate that drinking sparkling water suppresses both subjective and cognitive fatigue during prolonged esports play. This effect is likely mediated by enhanced prefrontal activity via TRP channel stimulation, as indicated by the attenuation of pupil diameter reduction.

Furthermore, the fatigue-reducing effects of sparkling water appears to support fair play behavior while maintaining in-game performance. Ultimately, sparkling water could offer a low-risk, sustainable alternative to caffeine- or sugar-based interventions for mitigating cognitive fatigue in modern digital life.

## Data availability

The data supporting the findings of this study are available from the corresponding author upon request.

## Author contribution

W.K., S.M., and T.M. conceived and designed the study. S.T. and T.M. recruited participants, collected the data, performed the analyses, and interpreted the results. S.T. drafted the manuscript. W.K., S.M., and T.M. critically revised the manuscript. All authors approved the final version of the manuscript.

---

**Supplementary Figure 1.**
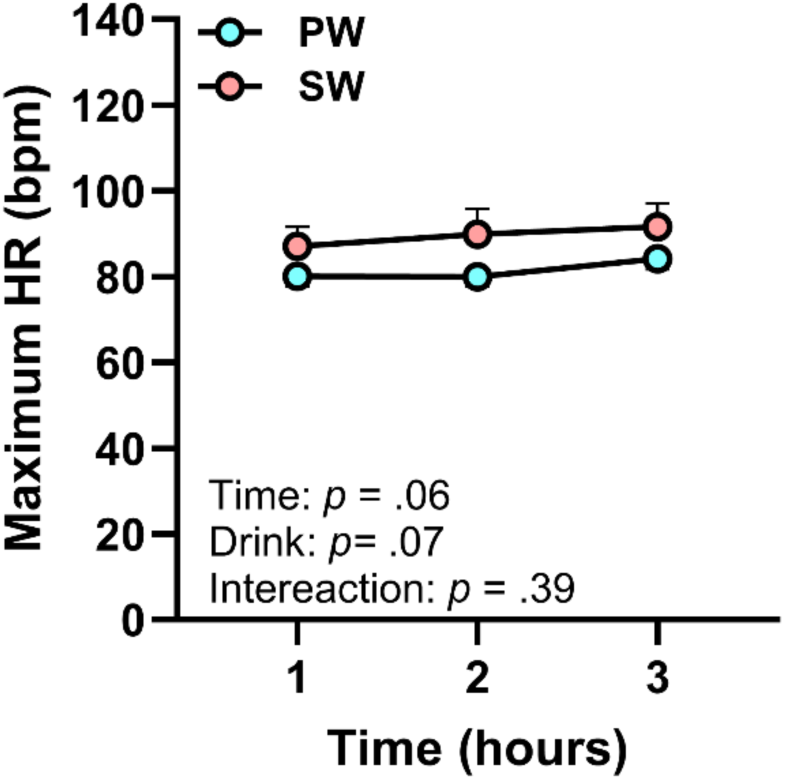
Drinking sparkling water does not increase average heart rate during esports play. Average hourly heart rate of participants during esports play. Data are shown as mean ± SEM. Results of two-way ANOVA are shown in the lower-left corner of each graph. *Time*: main effect of playing time. *Drink*: main effect of drink condition. *Interaction*: interaction between playing time and drink condition.

**Supplementary Figure 2.**
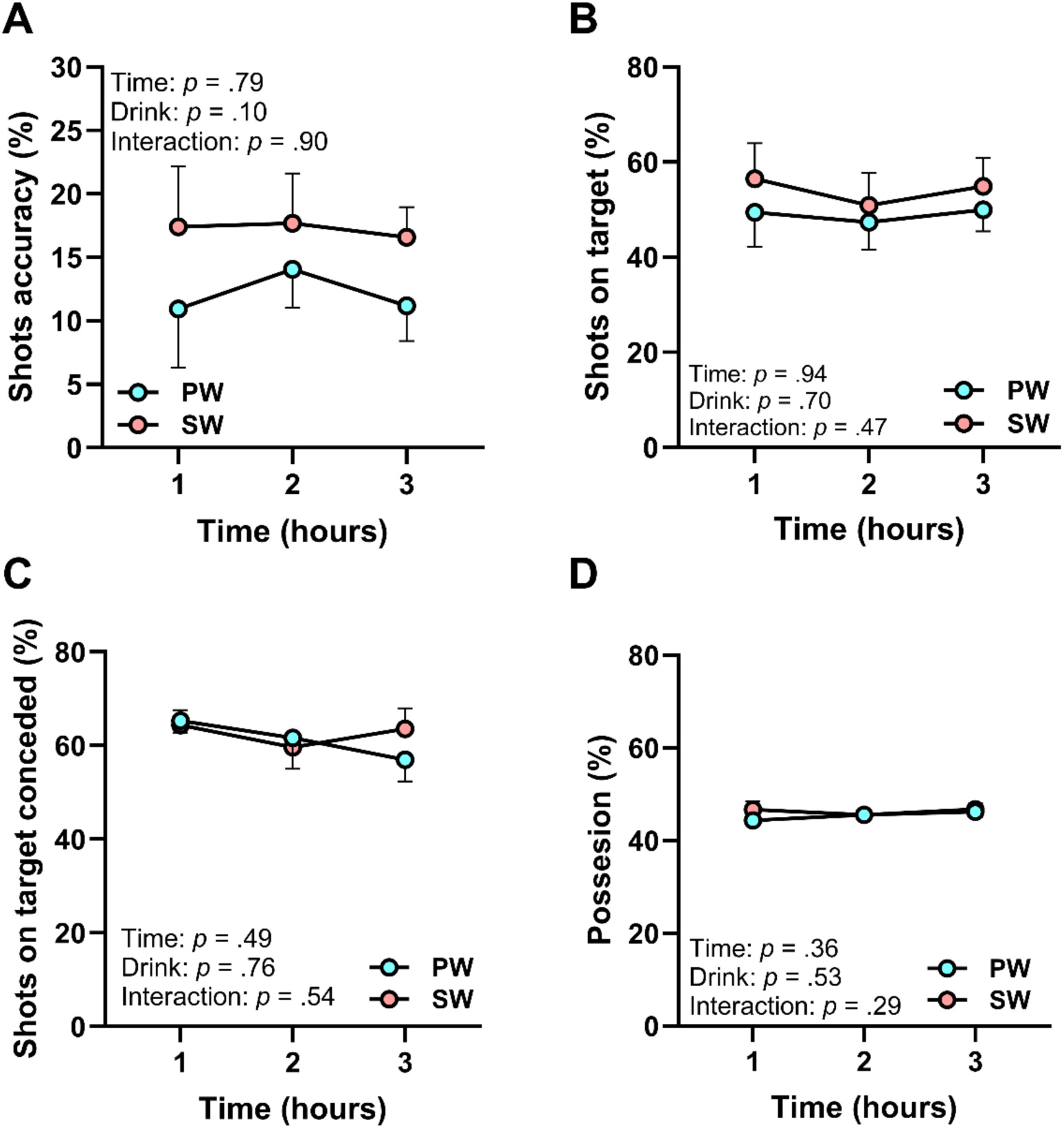
Drinking sparkling water does not affect offensive performances. **A.** The average number of goals. **B.** The average number of lost scores. **C.** The average number of shots. **D.** The average accuracy of shots. **E.** The average number of passes. **F.** The average number of intercepts. Data are shown as mean ± SEM. Results of two-way ANOVA are shown in the lower corner of each graph. *Time*: main effect of playing time. *Drink*: main effect of drink condition. *Interaction*: interaction between playing time and drink condition.

